# Tissue distribution of ACE2 protein in Syrian golden hamster (*Mesocricetus auratus*) and its possible implications in SARS-CoV-2 related studies

**DOI:** 10.1101/2020.06.29.177154

**Authors:** Voddu Suresh, Deepti Parida, Aliva P. Minz, Shantibhusan Senapati

**Affiliations:** Tumor Microenvironment and Animal Models Lab, Institute of Life Sciences, Bhubaneswar, India; Regional Centre for Biotechnology, Faridabad, Haryana, India

## Abstract

Recently, the Syrian golden hamster (*Mesocricetus auratus*) has been demonstrated as a clinically relevant animal model for SARS-CoV-2 infection. However, lack of knowledge about the tissue-specific expression pattern of various proteins in these animals and the unavailability of reagents like antibodies against this species hampers optimal use of these models. The major objective of our current study was to analyze the tissue-specific expression pattern of angiotensin□converting enzyme 2 (ACE2), a proven functional receptor for SARS-CoV-2 in different organs of the hamster. We have adapted immunoblot analysis, immunohistochemistry, and immunofluorescence analysis techniques to evaluate the ACE2 expression pattern in different tissues of the Syrian golden hamster. We found that kidney, small intestine, esophagus, tongue, brain, and liver express ACE2. Epithelium of proximal tubules of kidney and surface epithelium of ileum expresses a very high amount of this protein. Surprisingly, analysis of stained tissue sections for ACE2 showed no detectable expression of ACE2 in the lung or tracheal epithelial cells. Similarly, all parts of the large intestine (caecum, colon, and rectum) were negative for ACE2 expression. Together, our findings corroborate some of the earlier reports related to ACE2 expression pattern in human tissues and also contradicts some others. We believe that the findings of this study will enable the appropriate use of the Syrian golden hamster to carryout SARS-CoV-2 related studies.

## Introduction

The current outbreak of COVID-19 (Corona Virus 2019) caused by the SARS-CoV-2 virus was declared as a pandemic on 11th March 2020. Till now it has succumbed around 10.26 million people worldwide and almost 504.7 thousand people have lost their lives due to this pandemic (till 29th June 2020). Therefore, there is an urgent need to study this viral disease transmission, pathogenesis, prevention, and treatment. In this regard, the role of clinically relevant experimental animal models is crucial. No single species of animal might be able to exactly recapitulate all the SARS-CoV-2 infection-related events in humans. However, the use of different animal models will help to address questions in a more reliable and clinically relevant manner. At the same time, the exploration of multiple species susceptible to this virus might also help to identify the natural reservoir and potential carriers of this pathogen.

Syrian golden hamsters (*Mesocricetus auratus*) being a permissive host to multiple other viruses and a recognized model for respiratory syndrome coronavirus (SARS CoV) infection, it has drawn immediate attention for COVID-19 related studies^1^. Studies have shown that the Syrian hamster infected by SARS-CoV-2 manifest various clinical signs of COVID-19 in human patients^2–4^. Moreover, the pathology of this disease in hamsters resembles humans. The model has also demonstrated high transmissibility of SARS-CoV-2 among close contact animals. In a recent study, Sia S. F. *et al*. used male Syrian golden hamsters at 4-5 weeks of age to check the potential use of hamster as a model of SARS-CoV-2 infection^3^. Analysis of tissue samples from the infected animals showed pathological changes in the respiratory tract and presence of SARS-CoV-2 N protein at bronchial epithelial cells, pneumocytes, and nasal epithelial cells. Duodenum epithelial cells also stained positive for viral N protein. No apparent histopathological changes were observed in brain, heart, liver, and kidney on 5 dpi^3^. Although no infectious virus was detected in the kidney, low copies of the viral genome were detected on 2 and 5 dpi^3^. In a similar type of study, Chan FJ *et al*. by using male and female Syrian hamsters of 6-10 weeks old identified tissue damage and presence of viral N protein at different parts of the respiratory tract (nasal turbinate, trachea and lungs)^2^. The viral N protein was abundantly present in bronchial epithelial cells, macrophages, type I and II pneumocytes. At 4 dpi, N protein expression was found all over the alveolar wall^2^. The histopathological analysis also showed tissue damage and/or inflammatory lesions at multiple extra-pulmonary organs (intestine, heart, spleen, bronchial, and mesenteric lymph nodes); however, N protein expression was only detected in the intestinal epithelial cells^2^. In the very recent past, Syrian hamster model of SARS-CoV-2 infection has been instrumental to establish that passive transfer of a neutralizing antibody (nAb) protects SARS-CoV-2 infection^5^.

Studies have clearly shown SARS-CoV-2 binds to human angiotensin-converting enzyme 2 (ACE2) expressed by its target cells and use it as a functional receptor to enter into cells^6, 7^. Hence, drugs that could inhibit the binding of viral proteins (S-protein) to the ACE2 expressed on the target cells are assumed to be potential therapeutics against COVID-19. A recent study has shown that human recombinant soluble ACE2 (hrsACE2) blocks early stage of SARS-CoV-2 infection^8^. We have also proposed the bioengineered probiotics expressing human ACE2 as a potential therapeutics against SARS-CoV-2 infection^9^. The alignment of ACE2 protein of different species has suggested that the S protein may interact more efficiently with Cricetidae ACE2 than murine ACE2^10^. *In silico* analysis also shows possible interaction between SARS-CoV-2 spike proteins with Syrian hamster ACE2^2^.

At the time of ongoing COVID-19 pandemic, in addition to the vaccine and antiviral development, attentions have been made to target host proteins for therapeutic purposes. As discussed above, the pharmaceutical modulation of ACE2 expression or inhibition of its interaction with SARS-CoV-2 spike protein for COVID-19 therapy is a matter of current investigation at different parts of the world^11^. In these efforts, animal models will be instrumental to check the efficacies and safety of potential drug candidates against COVID-19. Although the Syrian hamster is a clinically relevant model for multiple infectious diseases, unavailability of reagents like antibodies against hamster proteins and lack of publicly available gene or protein expression data for this species are the major constrains to use these models up to their full capacity^12^. Before utilizing hamster as a model to understand the role of ACE2 in the pathogenesis of SARS-CoV-2 infection and/or to evaluate the efficacy of ACE2-targeted drugs, the knowledge about the basal level of ACE2 expression in different tissues of hamster is very essential. In the current study, we have checked the expression pattern of ACE2 in different tissues of normal Syrian hamster through immunoblot and immunohistochemical analysis.

## Material and Methods

### Isolation of hamster tissue samples

All the tissue samples used in this study are from archived samples collected during our previous studies^12, 13^. Prior approval from the Institutional Animal Ethical Committee (Institute of Life Sciences, Bhubaneswar, India) was taken for use of these animals. All the methods associated with animal studies were performed according to the Committee for the Purpose of Control and Supervision of Experiments on Animal (CPCSEA), India guidelines.

### Western blot analysis

Using an electric homogenizer, tissues were lysed in ice-cold RIPA buffer (20 mM Tris-HCl pH 7.5, 150 mM NaCl, 1 mM Na2 EDTA, 1 mM EGTA, 1% NP-40, 1% sodium de-oxy-cholate, 2.5 mM sodium pyrophosphate, 1 mM β-glycerophosphate, 1 mM Na3VO4) supplemented with a protease inhibitor cocktail (MP Biomedicals) and soluble proteins were collected. Protein concentrations were measured by Bradford assay (Sigma). 20 μg of protein was loaded for each sample and electrophoresed through 8% SDS-polyacrylamide gels. Proteins were transferred to poly-vinylidene difluoride membrane (Millipore) and blocked with 5% bovine serum albumin. Membranes were probed with ACE2 (#MA5-32307; Invitrogen; 1:3000) or β-actin (#A2066; Sigma-Aldrich; 1:1000) primary antibody and horseradish peroxidase-conjugated secondary antibody. Antibody binding was detected with electrochemiluminescence substrate (#12757P; CST) and chemiluminescence visualized with ChemiDoc™MP Gel Imaging System (BioRad).

### Immunohistochemistry (IHC)

All the tissue samples were processed and sectioned as reported earlier ^12, 13^. Paraffin-embedded sections were de-paraffinized using xylene, rehydrated in graded ethanol and deionized water. Sections were subjected to antigen retrieval treatment by boiling in acidic pH citrate buffer (Vector Laboratories) for 20 min in a steam cooker. 3% hydrogen peroxide in methanol was used to block the endogenous peroxidase for 20 min and washed with 1X PBS two times followed by blocking with horse serum (Vector Lab) for 30 min at room temperature. Sections were treated with ACE2 antibody (#MA5-32307, Invitrogen, 1:200) overnight in a humidified chamber at 4 °C. Sections were washed twice with 1X PBS for 5 min each. Slides were treated with horse anti-rabbit/mouse IgG biotinylated universal antibody (Vector Laboratories) for 45 min at room temperature and with ABC reagent for 30 min. To develop the stain 3, 3’-diaminobenzidine (DAB; Vector Laboratories) was used as a substrate according to the manufacturer’s instructions and hematoxylin was used as a counter-stain. Sections were dehydrated with ethanol, cleared with xylene, and mounted with Vecta mount permanent mounting medium. Sections were observed under microscope (Leica ICC500) and images were captured at 40x magnification.

### Immunofluorescence

Paraffin sections were subjected to de-paraffinization, rehydration, and antigen retrieval treatment. Sections were blocked with horse serum (Vector Lab) for 30 min at room temperature and probed with ACE2 primary antibody (1:200) overnight in a humidified chamber at 4°C. Sections were washed twice with PBST and treated with anti-Rabbit Alexa Fluor 594 (#A-11037; Life technologies, 1:500) at room temperature for 45 min under dark conditions. After washing, slides were mounted with SlowFade Gold Antifade mountant with DAPI (#S36938; Life Technologies) and visualized using TCS SP8STED confocal microscope.

## Results and discussions

The ACE2 recombinant rabbit monoclonal antibody (Thermo Fisher Scientific; clone SN0754; Cat No MA5-32307) used in this study was generated by using synthetic peptide within human ACE2 aa 200-230 as immunogen. As per the information available by different companies, the antibody of this clone (SN0754) has reactivity against human, mouse, rat, and hamster (Thermo Fisher Scientific: MA5-32307 and Novus Biologicals: NBP2-67692). There are 26 species of hamsters in the world. In the recent past Syrian golden hamster (*Mesocricetus auratus*) has been demonstrated as a clinically relevant model for SARS-CoV-2 infection^2–4^. Hence, to check whether this antibody has reactivity against Syrian golden hamster ACE2, we initially did an immunoblot analysis of proteins isolated from different tissues of these animals. The antibody showed clear reactivity with a protein of molecular weight of ~ 120 kd, which matches with ACE2 (**Figure 1**). The absence of any other non-specific bands in the western blot also suggests its suitability for use in IHC of different organs of hamsters.

**Figure 1:**
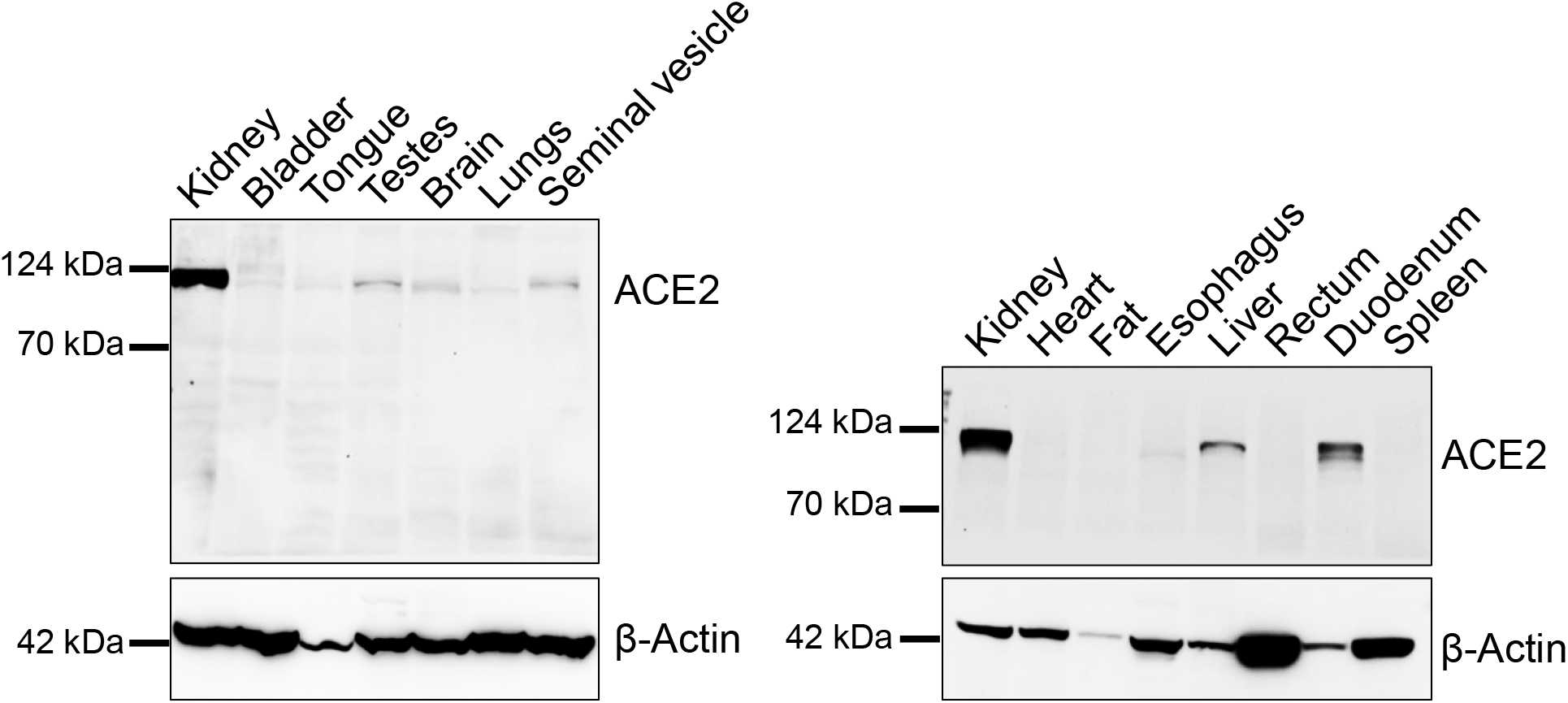
Expression status of ACE2 in different organs of Syrian golden hamster. Immunoblot analysis showing the expression status of ACE2 in tissues of multiple organs of the Syrian golden hamster. β-actin was used as an internal control.

The major objective of this study was to check the status of ACE2 expression in different hamster tissues. In human patients, the lung associated pathology is a predominant feature of SARS-CoV-2 infection^14^. Certain earlier studies have shown expression of ACE2 transcripts or protein by lung epithelial cells^15–18^, hence, just after the reports that ACE2 binds with SARS-CoV-2 spike (S) protein, the research and clinical communities assumed that high level of ACE2 expression in lung or other part of the respiratory tract might be a major driving factor in the pathogenesis of this respiratory virus. Our initial immunoblot analysis data showed very trace amount of ACE2 expression in lung tissue lysate (**Figure 1**). To get an idea about spatial and cell-type distribution of ACE2 expressing cells in lungs and trachea further IHC analysis was conducted. Interestingly, we didn’t find any visible positive staining in the epithelial cells of trachea, bronchioles, and alveoli (**Figure 2A**). Endothelial cells and smooth muscle cells associated with the wall of blood vessels were also negative for ACE2 staining (data not shown). As most of the previous reports suggest ACE2 expression in type II pneumocytes, we performed immunofluorescence staining to get magnified images for alveolar pneumocytes and tracheal epithelial cells. Corroborating our IHC data, we didn’t notice any positive staining when compared with corresponding without antibody stained tissue sections (**Figure 2B**). We believe that the trace amount of lung-associated ACE2 detected in immunoblot analysis might have come from some non-epithelial cells with low abundance and scattered distribution in lung parenchyma, whose presence might not be obvious in stained tissue sections. Our findings corroborate a recent report on human ACE2 expression patterns in different organs published in a preprint form^19^. Together, based on our preliminary findings and available reports^19^ we believe that SARS-CoV-2 related lung pathology might be independent or minimally dependent on ACE2 expression status in lungs, which warrants further investigation. Recent studies have reported the presence of SARS CoV-2 viral protein in respiratory tract epithelial cells and lungs of infected hamsters^3^, hence our findings suggest the possible involvement of some other proteins than ACE2 in entry of SARS-CoV-2 virus into respiratory and lungs cells^20, 21^.

**Figure 2:**
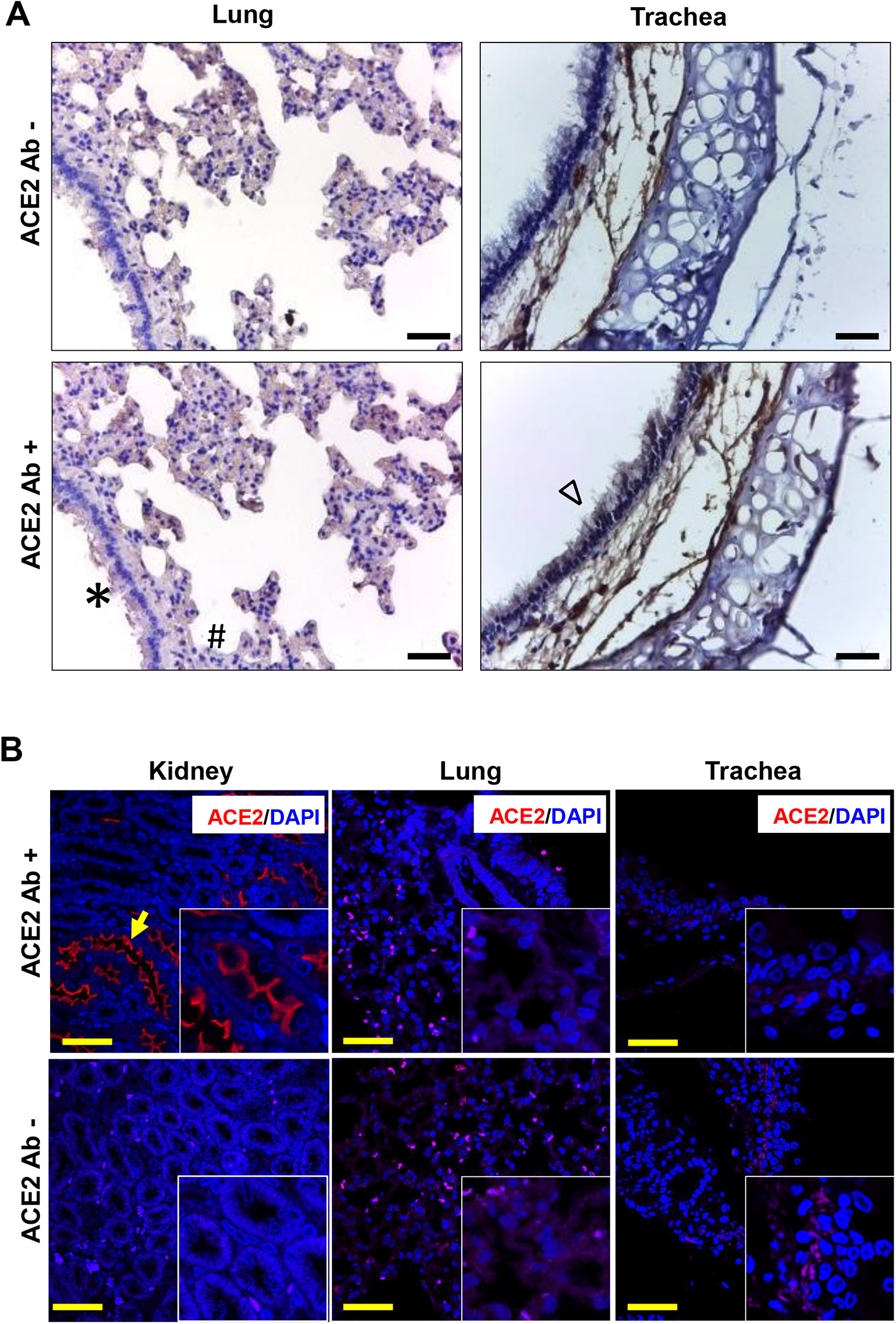
ACE2 expression pattern in hamster lung and trachea tissues. **(A)** Micrographs showing immunohistochemical staining of hamster lung and trachea tissues for ACE2. Corresponding tissues stained without primary antibody were used as negative controls. Cells marked with *, # and Δ symbols indicate bronchial, alveolar, and tracheal epithelial cells (scale bar =50μm). **(B)** Micrographs showing immunofluorescence staining of hamster lung, trachea, and kidney tissues with ACE2 primary antibody. Corresponding tissues stained without primary antibody were used as negative controls. The yellow arrow symbol indicates positively stained cells (scale bar = 25μm).

In our study, hamster kidney tissues showed very high level of ACE2 expression (**Figures 1, 2B & 3**). Its expression was mostly at the apical surface of proximal tubules whereas glomeruli were negative (**Figures 2B & 3**). So far most of the literature and publicly available protein expression database have clearly shown high expression of ACE2 protein in human kidney tissues^8, 22^. High expression of ACE2 in the kidney is believed to contribute to SARS-CoV-2 virus pathogenesis and disease severity^22^. Detection of kidney injuries in tissues of COVID-19 patients’ post-mortem tissues further supports the importance of considering kidney function-related issues for COVID-19 treatment and management^23^. Detection of SARS-CoV-2 viral genome in certain patients’ urine samples has also been reported^24^. Using a kidney organoid model, Monteil V *et al*. have demonstrated that the proximal tubules express ACE2 and SARS-CoV-2 replicated in these organoids^8^. Despite this clinical and experimental evidence from human patients or tissues, so far none of SARS-CoV-2-related studies in hamsters have reported any kidney-related histopathological changes^2–4^. In the future, further investigation and analysis are required to confirm whether hamster kidney epithelial cells are permissible to this viral infection. The tropism of virus to different organs depends on multiple factors including the organ-specific microenvironment. At the same time, infectivity of isolates from different parts of the world is yet to be experimentally compared. In future, studies with different isolates and/or hamsters with different comorbidity conditions might help to decipher the role of kidney cells in SARS-CoV-2 pathogenesis.

**Figure 3:**
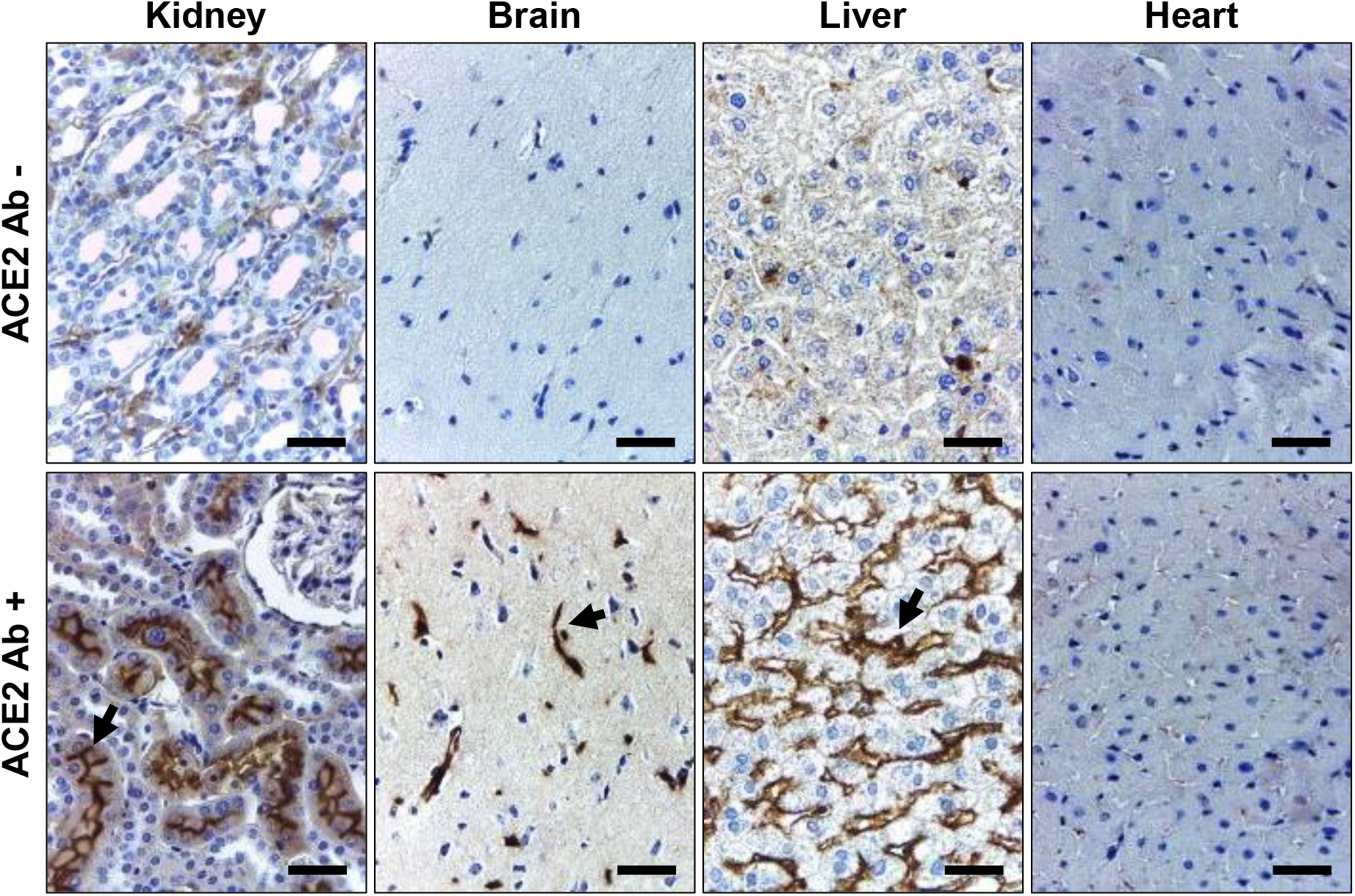
ACE2 expression pattern in hamster vital organs tissues other than lungs. Micrographs showing immunohistochemical staining of hamster vital organs other than lungs for ACE2. Corresponding tissues stained without primary antibody were used as negative controls. The arrow symbol indicates positively stained cells in different tissues (scale bar =50μm).

Like ACE2 expression in kidney tissues, consistently we observed a high level of ACE2 expression at different parts of the hamster gut (**Figure 4**). Prior studies have reported high expression of ACE2 in human gut tissues^16, 19^. Single-cell transcriptomic analysis of gut tissues has identified expression of ACE2 in upper epithelial and gland cells of esophagus and absorptive enterocytes of ileum and colon^18^. Our IHC data shows high expression of ACE2 in the surface epithelial cells of hamster esophagus, duodenum, ileum but no expression in the large intestine (caecum, colon, and rectum) (**Figure 4**). ACE2 expression in enterocytes indicates the possibilities for these cells being infected by SARS-CoV-2 through intestinal routes. Our study and from other parts of the world have clearly shown shedding of viral genome in COVID-19 patients stool samples^25^. Importantly, studies have also reported isolation of infectious SARS-CoV-2 virus from the stool samples^26^. In the recent past, some elegant studies have clearly shown infection of gut epithelial cells by SARS-CoV-2^27, 28^. Diarrhoea is a common finding in multiple COVID-19 patients across the world^29^ and in certain COVID-19 patients, gut epithelial cells damage has also been reported. Hamster model with SARS-CoV-2 infection also showed intestinal epithelial damage and expression of viral protein in enterocytes^2^. These data suggest that ACE2 expressed by gut epithelial cells might have a role in the pathogenies of SARS-CoV-2 infection. Considering these points, hamster might be a suitable model to investigate the intestinal pathogenesis of SARS-CoV-2 infection and evaluate different therapeutics that target ACE2.

**Figure 4:**
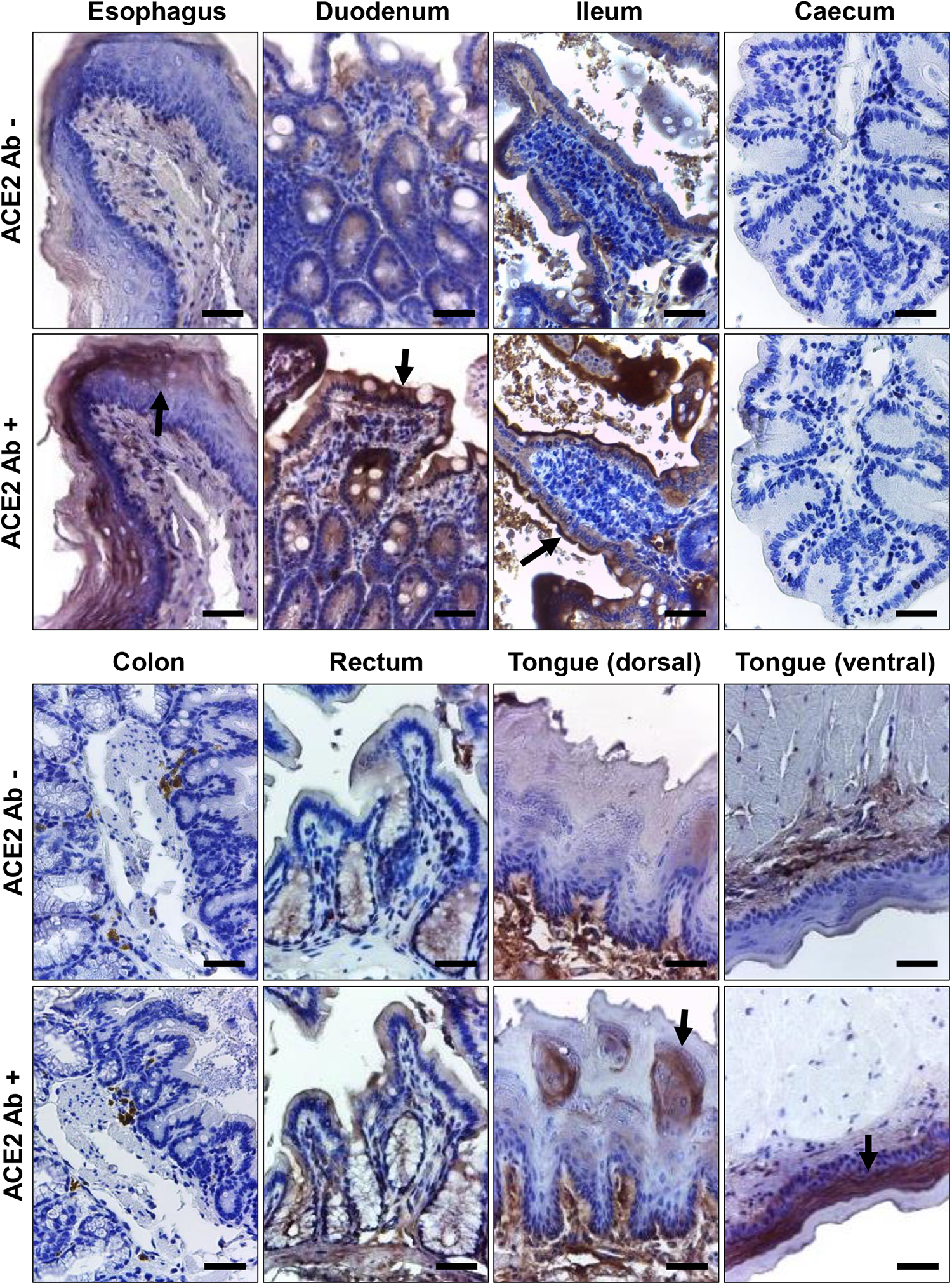
ACE2 expression pattern in different parts of hamster gastrointestinal tract and tongue tissues. Immunohistochemical analysis showing representative images of ACE2 expression at different parts of hamster gastrointestinal tract and tongue tissues. Corresponding tissues stained without primary antibody were used as negative controls. The arrow symbol indicates positively stained cells in different tissues (scale bar =50μm).

In our analysis, in addition to kidney and different parts of the gut, brain, liver, tongue are three other extra-pulmonary organs that showed ACE2 expression (**Figure 1, 3 & 4**). In brain, mostly the neurons of the cerebral cortex were positive for ACE2 expression. This finding corroborates information available at the human protein atlas (https://www.proteinatlas.org/ENSG00000130234-ACE2/tissue). Expression of ACE2 in hamster brain neuronal cells might help in investing the possible neurological tissue damage due to SARS-CoV-2 infection^30^. In liver tissues, mostly the sinusoidal endothelial cells stained positive for ACE2, but hepatocytes were negatively stained (**Figure 3**). Our immunoblot analysis data also showed ACE2 expression in hamster liver tissues (**Figure 1**). The sinusoidal endothelial expression of ACE2 in hamster doesn’t match with the expression of ACE2 pattern reported for human liver^16^, and warrants further investigation to understand these contradictory findings. In our analysis, we didn’t notice any positive staining in bile duct and gall bladder epithelial cells (data not shown). The human oral mucosal cavity expresses ACE2 and specifically this is highly enriched in tongue epithelial cells^31^. The data from our IHC study also shows expression of ACE2 in both dorsal and ventral stratified squamous epithelium of hamster tongue. Interestingly, the ventral side epithelial cells have very high level of ACE2 expression than dorsal side (**Figure 4**). The absence of ACE2 expression in our immunoblot analysis (**Figure 1**) could be due to less proportion of cellular proteins contribution into the total tissue lysate (also a possible reason for low level of β-actin detection).

Together, our study has provided a comprehensive idea about ACE2 expression patterns in different tissues of hamster. We believe that this information will be instrumental in optimal use of the Syrian golden hamster as a model for SARS-CoV-2 infection. At the same time, whether expression of ACE2 in hamsters depends on age, sex or other pathophysiological conditions warrants further investigation. In the future, investigators might use different other antibodies to check the expression pattern of ACE2 proteins in hamster tissues, this information will further help in confirming our findings and more importantly to understand the possible effect of tissue-specific post-translation modification of ACE2 on antibody reactivity.

**Figure 5:**
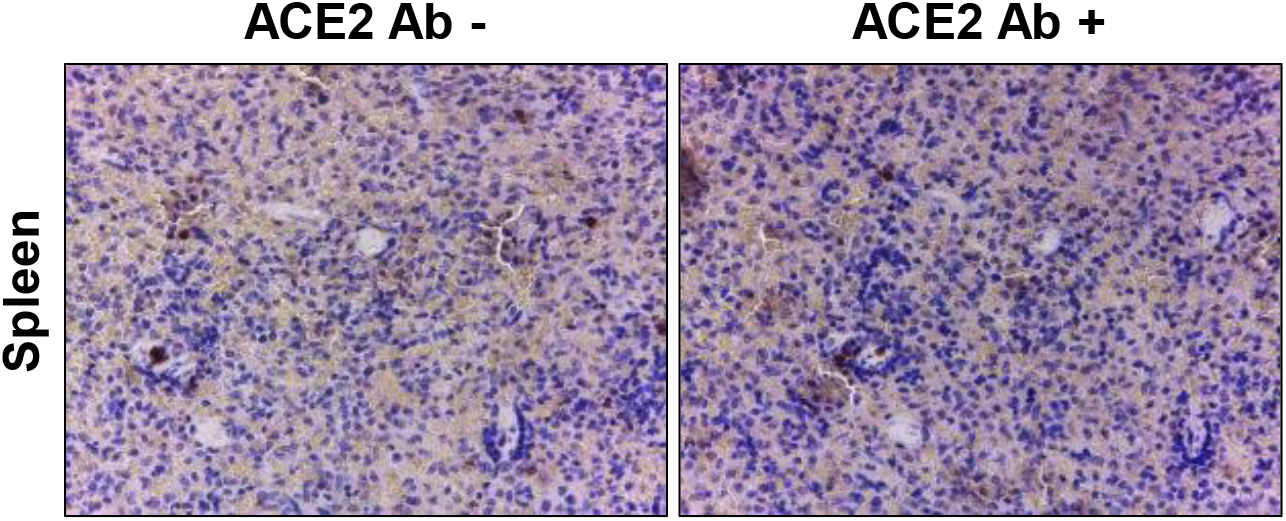
ACE2 expression status in hamster spleen tissues. Micrographs showing immunohistochemical staining of hamster spleen tissues for ACE2. Corresponding tissues stained without primary antibody were used as negative controls (scale bar =50μm)

## Acknowledgments

The authors are thankful to Director, ILS, Bhubaneswar for his support. VS, DP and APM are recipients of Council of Scientific and Industrial Research (CSIR) students’ research fellowship, Government of India. We sincerely acknowledge the technical supports given by Mr. Madan Mohan Mallick and Mr. Bhabani Sahoo, ILS.

